# Cognitive effort increases the intensity of rewards

**DOI:** 10.1101/2023.08.24.554617

**Authors:** Mejda Wahab, Nicole L. Mead, Stevenson Desmercieres, Virginie Lardeux, Emilie Dugast, Roy F. Baumeister, Marcello Solinas

## Abstract

An important body of literature suggests that exerting intense cognitive effort causes mental fatigue and can lead to unhealthy behaviors such as indulging in high-calorie food and taking drugs. Whereas this effect has been mostly explained in terms of weakening cognitive control, cognitive effort may also bias behavioral choices by amplifying the hedonic and emotional impact of rewards. We report parallel findings with animals and humans supporting this hypothesis. In rats, exerting cognitive effort immediately before access to cocaine self-administration significantly increased drug intake. In addition, exerting cognitive effort increased the psychostimulant effect of cocaine. The effects of cognitive effort on addiction-related behaviors were eliminated and even reversed when animals could rest in their home-cage for 2-4h before access to cocaine self-administration. Among humans, we found that expending cognitive effort increased consumption of tasty (but unhealthy) food by increasing the hedonic enjoyment of consuming the food. In addition, the effects were specific for emotionally relevant stimuli (i.e., food rewards) and did not generalize to judgment about neutral objects. Altogether these data suggest that intense cognitive effort can increase the perceived intensity of rewards and lead to their overconsumption. This effect may contribute to bad decision making induced by excessive cognitive effort and make people more vulnerable to indulge in unhealthy behaviors such as use of addictive drugs.

**Significance Statement:** People dieting or recovering from addiction frequently report that relapses occur during periods of stress and mental fatigue. Multiple processes may contribute to this, including beliefs about the stress-reducing effects of drugs, beliefs about one’s inability to continue resisting, and lack of energy needed to sustain resistance. Here, we suggest an additional possible process: during a state of mental fatigue, rewards become all the more satisfying, thereby also increasing subsequent desire for them. We report two lines of experiments, one with rats and one with human participants, to show that intense cognitive effort leads to increased enjoyment of rewards and hence increased consumption.

## Introduction

Making decisions and regulating one’s own responses are essential for adaptive behavior. Evidence suggests that the human capacity for decision making and self-regulation is limited and that following effortful acts of choice or self-control, people have reduced capacity to exert inhibitory control (1, 2). Thus after expending cognitive effort, people are more likely to engage in unhealthy activities such as overeating (3, 4) and using drugs (5, 6). Whereas there have been some controversies about the replicability of these findings (7), recent metanalyses show that self-control failure (also known as ego depletion) is one of the most solid findings in social psychology (8), especially when the duration and intensity of cognitive effort are high.

Many behavioral conflicts can be conceptualized as a balance between impulse and restraint (9). Behavioral disorders such as addiction are associated with imbalances between subcortical Go signals and cortical no-Go signals (10). Whereas the self-control failure has been mostly assumed to depend on weakening of restraint while leaving the impulses unchanged, it has recently been suggested that cognitive effort may intensify rewards, by making people feel them more intensely, which could contribute to unhealthy choices (11). Put simply, after an effort, individuals can be more prone than others to yield to temptation, either because their resistance is weaker or because the temptations are felt more strongly (or both). Whereas scattered evidence suggests both may be correct (11), experimental demonstration for intensified desire is still mostly lacking.

In this investigation, we performed experiments on both laboratory animals and humans that show that cognitive effort intensifies the rewarding effects of drugs and palatable food. The rat study manipulated effort by requiring rats to adjust their behavior repeatedly to changing contingencies within a complex operant task (12, 13). To rule out non-specific effects of operant training and food consumption during the cognitive training sessions, control rats did not have to adjust their behavior and received the same rewards easily. Next, some rats were given the opportunity to self-administer either cocaine or saline solution. Others rested for 2-4 hours and then performed the self-administration exercise. In addition, in separate groups of rats, we tested whether cocaine-induced locomotor effects, which are directly related to cocaine’s ability to activate the dopamine reward system (14), were altered by cognitive effort. If cognitive effort and resulting fatigue increase reward sensitivity, then rats should self-administer the most cocaine when doing so immediately after the effortful task (as compared with both the easy task and with the effortful task followed by restful delay) and cocaine-induced locomotor activity should increase.

To test our hypotheses with humans, we manipulated cognitive effort by having some participants (but not others) suppress a forbidden thought (i.e., a white bear; (15)) during a thought listing task. After this, participants self-administered potato chips under the guise of rating their taste and texture. Then they rated how much they enjoyed eating the chips. We predicted that expending effort on the cognitive task would increase hedonic enjoyment of the tasty but unhealthy food which in turn would lead them to consume more. In a follow-up study, we manipulated cognitive effort by having participants write a brief essay about their daily routine while avoiding using either two common letters (A and N) or two uncommon letters (X and Z) (16). We then tested the hypothesis that cognitive effort specifically heightens people’s desire for rewards rather than broadly increasing ratings of intensity, in this case perceptual judgments about the sensory qualities of neutral objects.

## Results

### Results in rats

#### Cognitive effort without rest increases cocaine intake

When cognitive and cocaine sessions were not temporally separated, performance in the cognitive sessions did not differ between the rats who later had cocaine vs. saline (Fig. S1, A-B). Even though the rate of success was different between cognitive effort and control groups (75% versus 100% of trials), the number of sugar pellets obtained in these sessions was similar in all groups (Fig. S2) indicating that controls waited longer in between trials.

When animals had performed the challenging cognitive exercise before they got access to the drug, they self-administered significantly more cocaine than control animals that had obtained the same food without making cognitive effort (Fig. 1). This was evident from a higher number of cocaine injections (Fig. 1A-B) in Coc-CE (cocaine and cognitive effort) compared to Coc-NoCE (no effort) rats. As expected, animals that were allowed to self-administer saline responded significantly less and self-administered fewer injections (Fig. 1) than both cocaine groups. Saline rats that performed the cognitive exercise (Sal-CE) did not differ from controls (Sal-NoCE) suggesting that the effects of cognitive effort are not simply due to non-specific effects of previous operant behavior or access to sweet food. Statistical analysis by three-way ANOVA on the number of injections per session revealed a significant effect of session [F (21, 903) = 8.11 ; p < 0.0001] demonstrating an increase over time, a significant effect of drug [F (1, 43) = 29.25; p < 0.0001] demonstrating more cocaine versus saline injections, a significant effect of cognitive exercise [F (1, 43) = 8.86; p=0.0048] demonstrating more injections in rats after a cognitive effort, a significant time X drug interaction [F (21, 903) = 1.77; p=0.018] showing that rats self-administering cocaine had higher increase over time than rats self-administering saline, and a significant drug X cognitive exercise interaction [F (1, 43) = 5.39; p=0.025] showing that cognitive effort increased cocaine self-administration more than saline self-administration.

**Figure 1.**
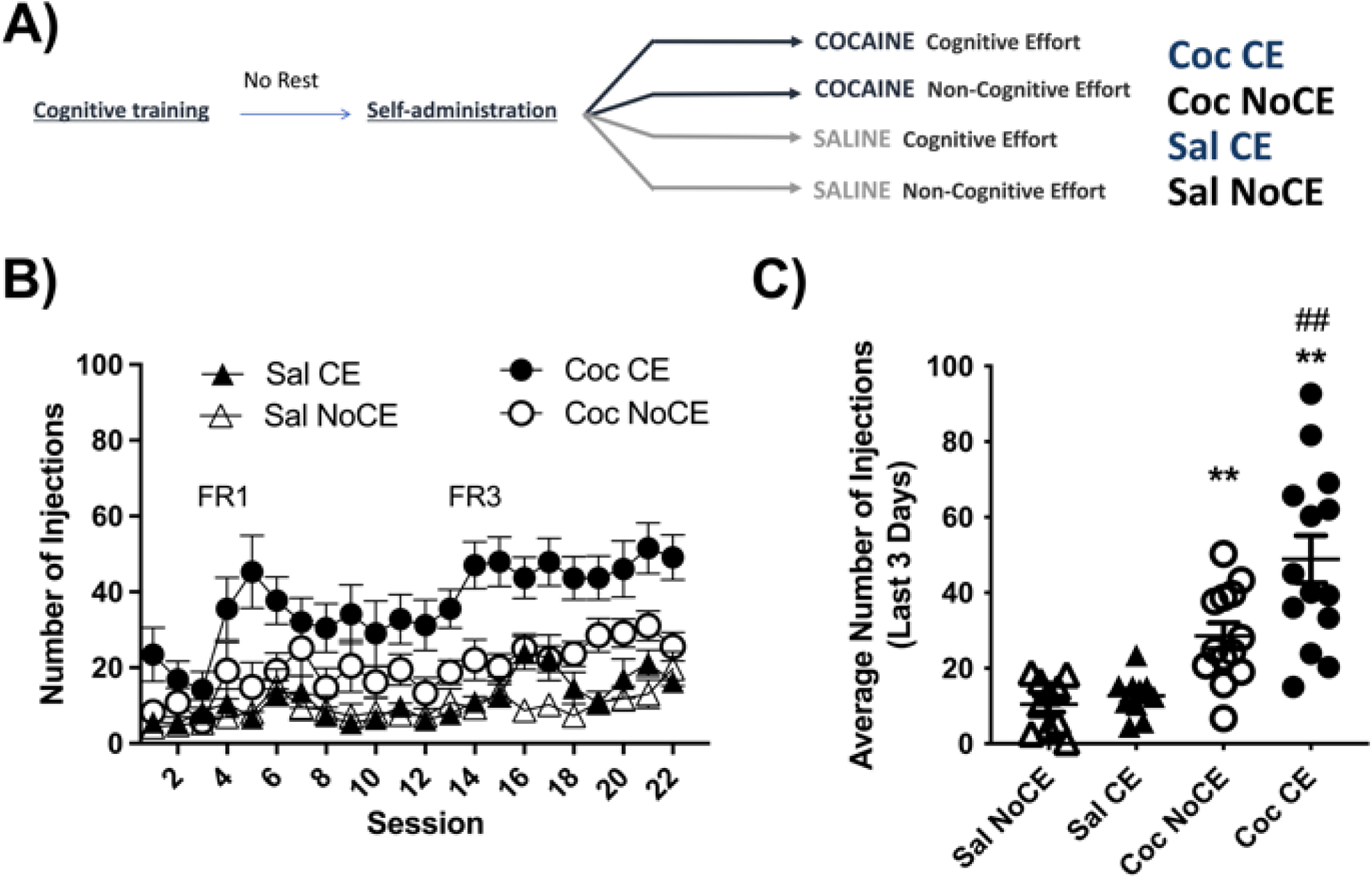
Effects of cognitive effort without rest on cocaine self-administration. Time course (A) and average in the last 3 days of self-administration (B) of number of injections in rats that performed a cognitive effort immediately before cocaine self-administration. Data are expressed as mean ± SEM (Sal NoCE, n =10; Sal CE n =10; Coc NoCE, n = 14; Coc CE, n = 13). Two-way ANOVA followed by post hoc Tukey’s test: **, P < 0.01 compared to Sal control; ##, P < 0.01 CE vs NoCE.

#### Rest after cognitive effort reduces cocaine intake

When cognitive and cocaine sessions were separated by 2-4h rest, performance in the cognitive sessions did not differ in Coc CE vs Sal CE rats (Fig. S1, C-D). On the other hand, the number of sugar pellets obtained in cognitive sessions was higher in control animals (that were always reinforced) compared to rats performing a cognitive exercise (that were reinforced on about 75% of trials) and this was regardless of whether the rats self-administered cocaine or saline (Fig. S3). Statistical analysis revealed a significant effect of cognitive exercise [F (1, 30) = 142.7); p < 0.0001] but no significant effect of drug or cognitive exercise X drug interaction.

After 2-4h rest, Coc-CE rats self-administered significantly less drug than Coc-No CE control rats (Fig. 2A-B). Again, saline rats self-administered a low number of injections, and no difference was found between Sal-CE and Sal-NoCE groups. Statistical analysis by three-way ANOVA of the number of injections per session revealed no significant effect of session [F (6.416, 192.5) = 0.6759; p =0.6793; Geisser-Greenhouse’s epsilon = 0.31] but a significant effect of drug [F (1, 30) = 28.20; p<0.0001] demonstrating that rats self-administered more cocaine than saline. There was also a significant effect of cognitive exercise [F (1, 30) = 6.587; P=0.0155] indicating that CE rats self-administered fewer injections than No CE rats, and a significant drug X cognitive exercise interaction [F (1, 30) = 7.80; p=0.0090] demonstrating that Coc-CE rats self-administered less cocaine than Coc-No CE rats.

**Figure 2.**
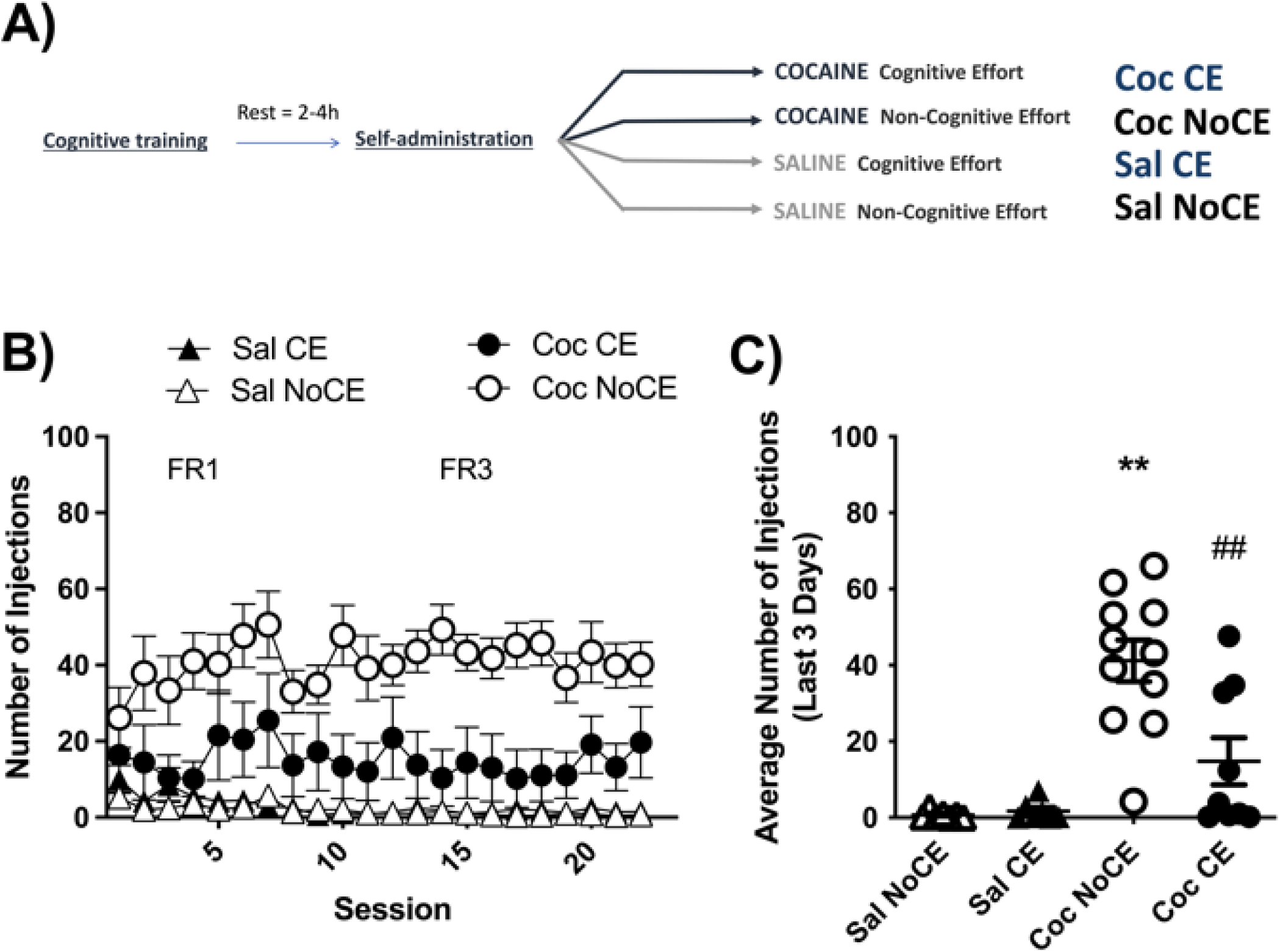
Effects of cognitive effort followed by 2-4h rest on cocaine self-administration. Time course (A) and average in the last 3 days of self-administration (B) of number of injections n rats that had a rest of 2-4h between cognitive exercise and cocaine self-administration. Data are expressed as mean ± SEM; (CE vs NoCE. Sal NoCE, n =7; Sal CE n = 7; Coc NoCE, n = 11; Coc CE, n = 9). Two-way ANOVA followed by post hoc Tukey’s test: **, P < 0.01 compared to Sal control; # and ##, P < 0.05 and 0.01 CE vs NoCE

#### Only cognitive effort without rest increases cocaine-induced locomotor activity

We then investigated, in separate groups of rats, the stimulant effects of passive injections of cocaine on locomotion, which are directly related to its ability to release dopamine (14) as a function of exerting cognitive effort, with or without rest. We found that compared to saline, cocaine increased locomotor activity in all groups (Fig. 3). However, rats that received cocaine immediately after a cognitive effort, but not rats that had rest between cognitive effort and the injection of cocaine, showed significantly more locomotor activity than all other groups (Fig. 3). Statistical analysis with one-way ANOVA demonstrated a significant effect of condition demonstrating that cognitive effort increases the stimulant effects of cocaine (F (4, 35) = 24.81; P < 0.0001).

**Figure 3.**
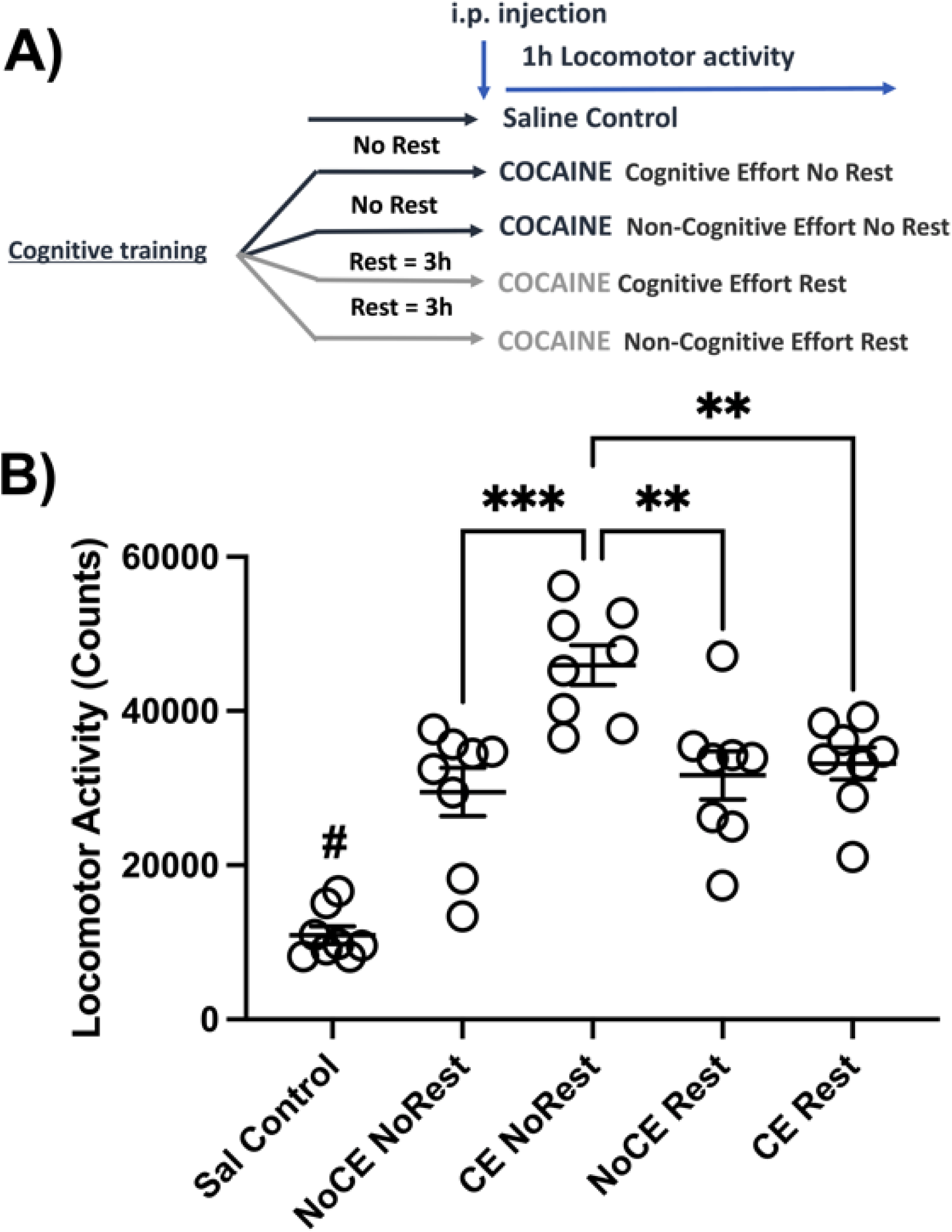
Effects of cognitive effort with or without rest on cocaine-induced locomotor activity. Locomotor activity measured for 60 min after injection of saline or cocaine (10 mg/kg, IP) in animals that performed a cognitive effort or control immediately before (0h) or 3h before the injection. (n =8 for all groups). One-way ANOVA followed by post hoc Tukey’s test: ** and ***, P < 0.01 and P < 0.001 different from CE Delay 0h; #, P < 0.0001 different from all other groups.

### Results in Humans

#### Cognitive effort increases enjoyment and consumption of tasty but unhealthy food

We report two experiments with human participants. The first investigated how cognitive effort affects the reward value and consumption of tasty but unhealthy food. Participants who had previously expended cognitive energy to suppress a forbidden thought during a thought listing task (vs. those who did not) reported enjoying the potato chips more, t(146) = 2.711, p = .008, d = .45 (M_high effort_ = 5.48, *SD* = 1.34; *M*_low effort_ = 4.85, *SD* = 1.45) (Fig. 4). Thus, expending cognitive effort intensified participants’ hedonic experience of consuming rewarding food.

**Figure 4.**
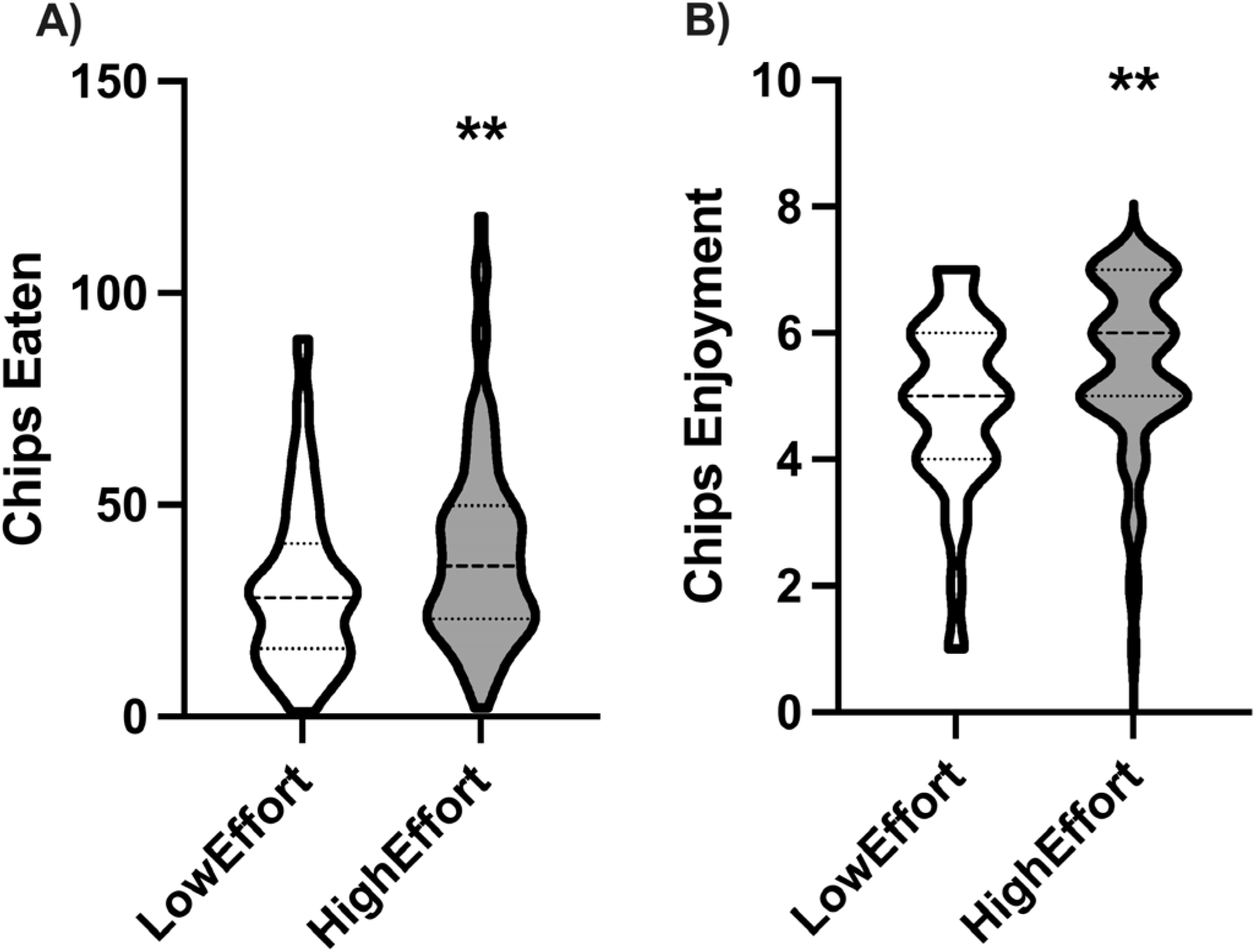
Effects of cognitive effort on enjoyment and consumption of chips. Violin plots show median and quartiles (High Effort, n = 80; Low Effort, n = 68). T-tests were used to compare the mean differences between the experimental conditions. * P < 0.05.

This heightened enjoyment in turn increased consumption. Participants who had previously expended cognitive effort consumed more potato chips than those who had not, t(146) = 2.475, p = .014, d = .41 (*M*_high effort_ = 39.56grams, *SD* = 23.49; *M*_low effort_ = 30.63, *SD* = 19.74) (Fig. 4). This translates into an increase of approximately 48 calories (from about 164 to 212). Amount eaten was significantly related to consumption enjoyment (*r*(147) = .25, *p* = .003; Fig. S4), so we conducted a bootstrapped mediation analysis (17). This model indicated that the effect of cognitive effort on increased eating was statistically explained by increased enjoyment (Fig. 5). Supporting the notion that cognitive effort increased consumption through heightened enjoyment, the direct effect of cognitive effort on consumption was rendered non-significant when chip enjoyment was included in the model (95% CI [-.307 to 14.013). More important, the indirect effect of cognitive effort on chip consumption via enjoyment was significant (95% CI [.362 to 4.410]).

**Figure 5.**
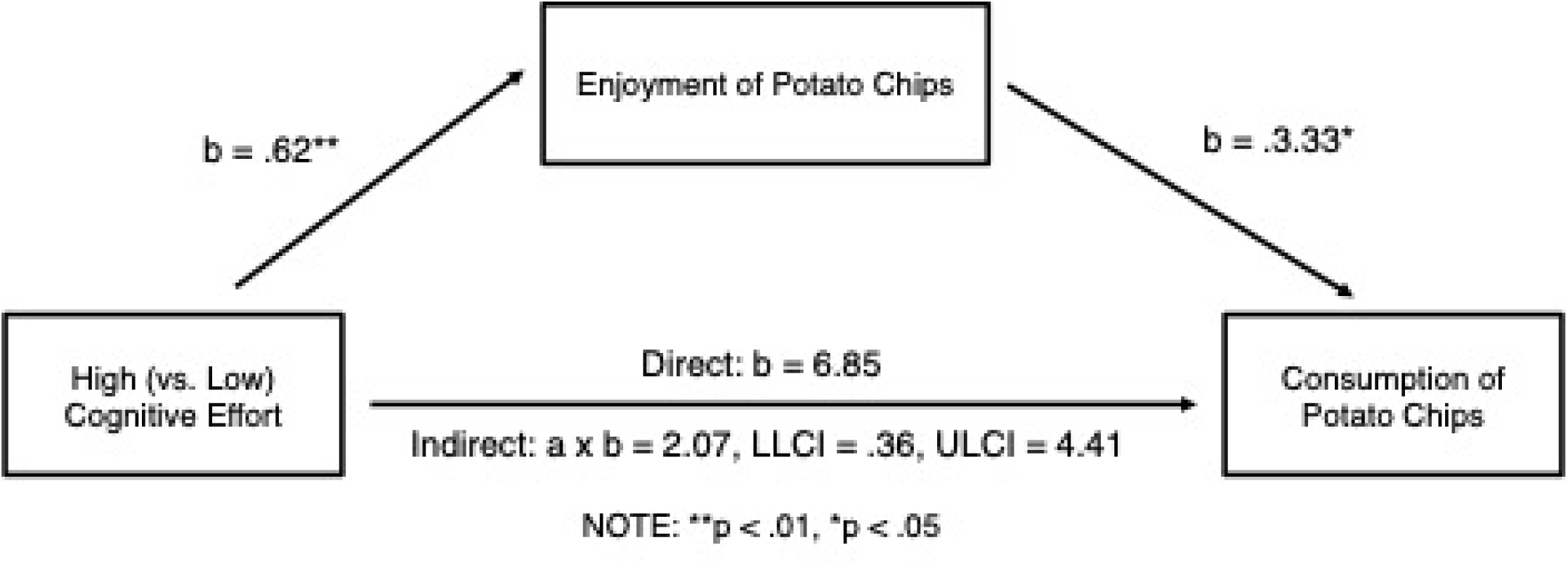
High (vs. low) cognitive effort increased potato chip consumption through heightened enjoyment of the potato chips. This figure displays the unstandardized effects obtained from process Model 4 (mediation; High Effort, n = 80; Low Effort, n = 68).

#### Cognitive effort affects perceptions of rewarding but not neutral objects

An alternative explanation could be that cognitive effort intensifies all ratings rather than just increasing sensitivity to rewards. To rule out this possibility, in a second experiment, we compared the effects of cognitive effort on perceptions of tasty yet unhealthy food and the sensory quality of neutral objects. After completing a writing task that did or did not require cognitive effort, participants tasted and then rated one chocolate. Then they rated a set of neutral office objects, such as how long was a pen or how bright was a yellow post-it note.

Consistent with expectations, expending cognitive effort intensified feelings toward chocolate specifically but not judgements about other objects (Fig. 6). A 2 (cognitive effort; no-cognitive effort) X 5 (type of item) repeated-measures ANOVA revealed the predicted interaction between cognitive effort and type of object, F (4, 95) = 2.967, p = 0.020, *h*_p_^2^ = 0.030. There was also a main effect of item type, F(4, 95) = 56.414, p < 0.001, *h*_p_^2^ = .373.

**Figure 6.**
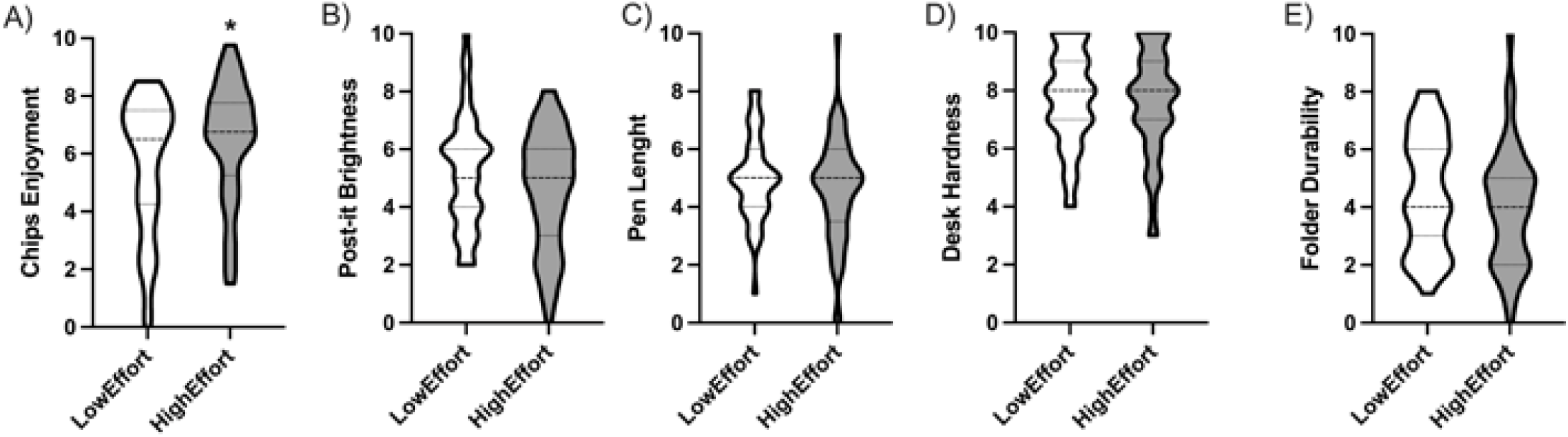
Effects of High vs Low effort on chocolate enjoyment and evaluation of neutral objects. Violin plots show median and quartiles (High Effort, n = 49; Low Effort, n = 48). T-tests were used to compare the mean differences between the experimental conditions. * P < 0.05.

Decomposing this interaction yielded results that were supportive of hypotheses. Participants who had previously exerted cognitive effort rated the chocolate as more enjoyable (M = 6.46, SD = 2.05) than participants who had not exerted a high amount of cognitive effort (M = 5.53, SD = 2.36) t(95) = 2.076, p = 0.041, d = .422. Follow-up analyses, broken down by chocolate question, revealed that participants who had completed the high (vs. low) cognitive effort task rated the chocolate as being more “yummy” (*M*_high effort_ = 7.06, *SD* = 1.76; *M*_low effort_ = 6.19, *SD* = 2.27; *t*(95) = 2.123, *p* = .036), more enjoyable to consume (*M*_high effort_ = 6.98, *SD* = 2.24; *M*_low effort_ = 6.02, *SD* = 2.65; *t*(95) = 1.927, *p* = .057), and reported a stronger desire to eat more chocolate at the present moment (*M*_high effort_ = 5.73, *SD* = 2.86; *M*_low effort_ = 4.50, *SD* = 3.45; *t*(95) = 1.921, *p* = .058). The cognitive effort manipulation did not affect how good the chocolate made participants feel (t < 1.29).

Also as expected, exerting cognitive effort did not alter participants’ perceptions of neutral, non-rewarding objects (ts < 1.44). These findings rule out the simple alternative explanation that the aftermath of cognitive effort simply makes one give more extreme ratings to everything. It did not change how people judged the length of a pen, the brightness of colored paper, or the hardness of their wooden desk. Its effects were specific to increasing desire for and enjoyment of rewarding stimuli.

## Discussion

Our studies with both rats and humans converged to show that the aftermath of cognitive, self-regulatory effort increases pleasurable desire and indulgence. After having to struggle to adapt to changing in contingencies, rats self-administered more cocaine and were more sensitive to the stimulant effects of cocaine than control condition rats. This was only found when cocaine was immediately available. Rats that were able to rest for a couple hours after the cognitive effort showed no increase in cocaine stemming from the cognitive exertion. In parallel, human participants consumed more chips following a cognitively demanding task than after an easier task, and heightened consumption of the tasty but unhealthy snack was caused by their increased enjoyment of consuming the chips. In a follow-up study, participants’ intensified hedonic desire after a more versus less cognitively demanding task was specific to a rewarding stimulus (chocolate) and was not found for neutral judgments about sensory characteristics of objects.

The notion that the capacity for self-regulation and other executive functions is limited is a staple of folk wisdom and more recently has received research support as well. Mental fatigue and declining ability to regulate attention, such as in vigilance tasks, have long been known in laboratory work (18). Research has shown with multiple methods and human populations that self-regulation deteriorates after exertion of cognitive effort. Although most theoretical accounts have focused on a transitory reduction in the ability to exert inhibitory control, recently it has been speculated that cognitive effort can also increase the value of rewards (11). Our results provide direct empirical support to this hypothesis. Thus, impulse and restraint would be causally intertwined. Just as the exertion of self-control leads to a weakening of restraint and control, it also causes feelings and impulses to be experienced more intensely than otherwise. Thus, when restraint is weak, desires and emotions are felt more strongly.

Addiction has always had a prominent place in self-control failure theory (18, 19). In fact, not only is addiction a disorder involving persistently bad decision making (20, 21), but it also implies 1) strong rewarding effects of drugs (22) and 2) need to exert cognitive inhibitory control of drug urges to limit drug consumption or to avoid relapse when abstinence is attained (10, 20). Our results suggest that all these processes can interact to drastically diminish the ability of patients to control their drug use. In fact, especially during periods of abstinence (23), resisting drug craving on the one hand would limit the ability to exert further inhibitory control and on the other hand it would amplify the rewarding effects of drugs.

Although we did not investigate the neurobiological mechanisms involved in effort-induced increases in reward intensity, the present effects are likely mediated by interaction between the prefrontal cortex and the nucleus accumbens. These regions play a central role in the balance between inhibitory control and impulses. Indeed, the prefrontal cortex is the main region responsible for cognitive control in both humans and rats (24). Conversely, the nucleus accumbens is the main region involved in reward and motivation (22, 25), emotion and impulses (26). In addition, it has been shown that when individuals successfully exert cognitive control the prefrontal cortex is activated and the nucleus accumbens is inhibited. However, when self-control fails the cortex is inhibited (27, 28), glutamate concentration and glutamate/glutamine diffusion increase in the lateral prefrontal cortex (29) and if individuals indulge in consumption of sweet food, the nucleus accumbens is activated (11, 28). The prefrontal cortex sends excitatory glutamate inputs to the nucleus accumbens (30) and interactions between glutamate and dopamine receptors in this brain region, is critical for the activating and reinforcing effects of cocaine (31, 32). Thus, it is possible that effort-induced decreases in prefrontal cortex activity alters glutamate neurotransmission in the nucleus accumbens leading to potentiation of dopamine signal and increased rewards.

Self-control failure has been mostly framed as a purely human phenomenon. Although there is some limited evidence of self-control failure outside of humans (33, 34), the human self-regulatory system is presumably highly advanced and atypical. Indeed, an explanation of self-control failure has been the constant need of humans to control their emotions in social settings to avoid conflicts (2). The present findings suggest that at least some effects of self-control failure occur in nonhuman animals, indicating an ancient and general process that is shared by humans and some mammals. A question arising from these findings is whether the effects of cognitive effort on reward intensity have an evolutionary role, or are they a byproduct of evolution? A possible role of effort-induced increase in reward is to reduce stress associated with cognitive effort. Indeed, whereas low to moderate levels of stress can be beneficial (“eustress”), prolonged, overwhelming stress can have negative effects on health and well-being (35). Thus, reducing the stress load can have beneficial effect for the individual. Furthermore, it has been shown that pleasurable behaviors such as consuming a sucrose solution decreases the neuroendocrine, cardiovascular, and behavioral responses to stress (36). In addition, stimulation of dopamine receptors (37) and cocaine, under certain circumstances (38), have been shown to produce anxiolytic effects. Thus, increasing the intensity of reward would oppose effort-induced stress effects and favor well-being. Importantly, whereas we have found evidence that effort-induced increases in emotional responses exist in both humans and animals, it remains to be determined whether the more commonly described effects on inhibitory control processes, which involve cortical functions that are clearly more developed in humans, also occurs in nonhuman animals.

Self-control exhaustion is believed to have a “refractory period during which time control is less likely to succeed” (1, 39), indicating that after an appropriate period of rest, self-control would be replenished and inhibition of inappropriate behavior would be effective again (1, 18, 40). Therefore, we expected that after appropriate rest, the effects of cognitive effort would recede and cocaine self-administration would not differ between Coc-CE and Coc-NoCE. We did indeed find that the effects receded. In fact, and somewhat surprisingly, when we imposed a delay between cognitive effort and availability of cocaine, self-administration of cocaine decreased even compared to controls. Although the self-control strength model also predicts that cognitive exercise, performed under appropriate conditions, should have beneficial effects on addiction (1, 18, 40), we believe that it is unlikely that these results are due to increased self-control for several reasons. First, as for experiments without rest, our procedure did not imply conflict and therefore, rats were not required to exert self-control. Second, in our locomotor activity experiments, the effects of cocaine after a cognitive effort followed by a period of rest, were similar to control rats that did not exert cognitive effort. Rats that rested after cognitive effort consumed less sucrose pellets than rats not performing a cognitive exercise but, since sucrose consumption has been shown to reduce cocaine intake (41), this would be expect to increase rather than reduce cocaine reinforcement. A possible explanation of these effects is that daily exposure to cognitive exercise acted like a form of environmental enrichment which has been consistently shown to reduce the rewarding effects of cocaine (42–44). However, these beneficial effects only occurs when cognitive exercise is followed by rest and are lost when cognitive effort is immediately followed by access to drugs.

This study has several limitations that should be acknowledged. First, whereas self-control failure has been interpreted as the result of mental fatigue, we cannot be sure that results in rats are due to real mental fatigue because this construct is difficult to extrapolate from rats’ studies. An alternative possibility is that rats performing in the cognitive task are simply more engaged in behavior and automatically respond for cocaine regardless of the reinforcing effects of cocaine. However, the control rats were at least as engaged in the operant task as cognitive effort rats so that nonspecific transfer of operant responding from food to cocaine can be ruled out. Another possibility is that increased cocaine self-administration is the result to frustration (45) due to partial reinforcement in cognitive effort rats (70% of reinforced trials) compared to 100% reinforcement in control rats. If this were the case, we should expect rats that make more errors and therefore receive less rewards per attempt to be more frustrated and self-administer more. However, there was no correlation between the performance in the cognitive task and cocaine intake (Fig. S5). Finally, it is possible that cognitive effort increases stress levels that are known to increase the reinforcing effects of cocaine. However, this limitation is only apparent as it is quite possible (even likely) that cognitive effort activates the stress/arousal systems (noradrenaline, crf, orexin, etc.) and that this activation participates in the negative effects of mental fatigue. Importantly, mental fatigue is a complex construct that has been proved tricky to evaluate even in humans as discrepancies between self-reports, behavioral outputs and brain activities have been shown (29). Future studies combining modern neuroscience approaches to monitor brain activity in behaving rats will be needed to draw more definitive conclusions determine whether the effects found in rats and the effects reported in humans represent the same neurobiobehavioral process.

Other limitations of the rat studies are that they were performed only on males and only using FR schedules. Importantly, significant differences exist in self-administration of drugs in male and female rats (46) highlighting the usefulness of performing experiments with both sexes in order to extrapolate to human conditions (47). Nevertheless, this limitation is compensated by our experiments with humans in which both men and women were tested and showed no differences in the effects of high cognitive effort on reward enjoyment and consumption (Experiment 1: sex X depletion interaction, Chips Eaten: p = 0.87; Chips Enjoyment: p = 0.27; Experiment 2: sex X depletion interaction, Chocolate Enjoyment: p = 0.92). In addition, the interpretation of FR schedule is not straightforward because increases in number of injections could be interpreted both as an increase or a decrease in the reinforcing effects of cocaine (48). However, our locomotor data show that the activating effects of cocaine are increased by cognitive effort without rest compared to no effort controls. This supports the interpretation of an increase in the rewarding effects of cocaine. Future studies in rats are needed to address all these questions and perform mechanistical studies to better characterize the neurobiological underpinnings of these effects.

Another limitation is that we used palatable, sweet (chocolate) and salty (chips), food as rewards in humans and cocaine in rats. Using cocaine in non-dependent humans raises important ethical issues and was unfeasible. Conversely, studying the rewarding effects of food in rats was unfeasible because rats may get confused between two tasks (the cognitive task and the reward task) that use the same reward (sucrose pellets). Second, to induce self-control failure, we used a word writing task in humans and an attentional set-shifting in rats. The latter task was chosen because it requires several cognitive functions such as attention, inhibition and flexibility and therefore, it is quite demanding (49). In addition, it has been shown to depend on the prefrontal cortex (12, 49), to produce stable behavioral performance (12, 13) and to predict excessive self-administration of methamphetamine in rats (13). Future studies will be required to determine whether performing tasks that imply specific cognitive functions such as memory, attention or behavioral inhibition produce similar effects on cocaine self-administration. In addition, it will be important to determine whether the effort-induced increase in the intensity of reward is specific to cocaine or occurs also for other drugs of abuse.

Self-regulation theory long has focused on the processes of response inhibition and other controls (2, 54). The present findings indicate that desires and emotions are felt more intensely when the capacity for self-regulation has been diminished by prior exertion. It may be a cruel irony that temptations become more intense just when one’s guard is down. These results have important implication not only for drug addiction but for a wide-range of behavior in which self-control failure effects can lead to unhealthy behaviors (3, 4). Recognizing the intertwined nature of these processes may help people come to manage motivated behaviors more effectively.

## Materials and Methods

Extended methods can be found in the supporting information (SI).

### Experiments in rats

#### Subjects

Seventy-five adult male Sprague-Dawley rats (Janvier Labs and Charles Rivers, France) experimentally naïve at the start of the study were used. All experiments were conducted in accordance with European Union directives (2010/63/EU) for the care of laboratory animals and approved by the local ethics committees (COMETHEA).

#### General Experimental Design

For the entire duration of the experiment, rats were food restricted at 85–90 percent of their weight. Rats were pre-trained for 20 daily sessions on the set-shifting procedure in order to learn the task and obtain a stable level of flexibility (Istin et al. 2017) before catheter implantation. After recovery, rats were divided into 4 experimental groups for each time conditions: cocaine cognitive effort (Coc CE), cocaine non-cognitive effort (Coc NoCE), saline cognitive effort (Sal CE), saline non-cognitive effort (Sal NoCE). Cognitive training and cocaine self-administration took place in the same experimental chambers which contained simultaneously operanda (levers and nosepoke) for food and drug that were remotely controlled by a PC to be active or inactive during the specific part of the session. For experiment 1, there was no interval between the cognitive session and the self-administration and the change from food to drug self-administration was associated simply by the switching off of the house light. For experiment 2, after the cognitive sessions, rats were removed from the self-administration cage and placed in their housing cage for a 2-4h period of rest before the self-administration session.

#### Cognitive exercise

Behavioral flexibility sessions lasted 45 min During these sessions, rats assigned to the CE conditions performed a modified version of an attentional set-shifting task in which animals must alternate between a visual rule (“follow the light to identify the active lever”) and an egocentric rule (“ignore the light position and stay either on the right or left lever”) (12, 13) in order to give the correct response and be rewarded with a food pellet.. Rules changed without specific signal, whenever the rat performed 10 consecutive correct responses in a given rule. Rats assigned to the NoCE condition, underwent a procedure that was similar to the attentional set-shifting task except that every lever press during trials was reinforced with a pellet delivery regardless of position of the lever and whether the light was on or off. Thus, this group did not need to exert a cognitive effort to obtain rewards.

#### Self-Administration Procedure

Rats were allowed to self-administer cocaine or saline for sessions that lasted 150 minutes, according to Fixed Ratio (FR) schedule of reinforcement using a single nose-poke as operandum. The start of the self-administration session was signaled only by the switching off of the house light and intermittent illumination of the nose-poke. Initially, FR value was set at 1 and, after 7 sessions, it was increased to 3.

#### Locomotor activity experiments

Separate groups of rats were trained for 20 sessions in the set-shifting task. After this training, half of the rats continued to perform the cognitive task (cognitive effort, CE) to obtain food pellets whether the other half, were reinforced regardless of the rule (No Cognitive effort, NoCE) for 5 additional sessions. After the last session, rats were injected with cocaine (10 mg/kg i.p.) either immediately or 3h after the cognitive session and were put in an open field where locomotor activity was recorded by a viewpoint videotracking system (www.viewpoint.fr) for 1h.

#### Experiments in humans

These studies received approval from the institutional review board at which they were conducted. Informed consent was obtained from all participants prior to commencing the study.

In experiment 1, 168 students at a Dutch university participated in exchange for a small monetary payment (5 euros in 2013). Twenty participants failed the attention check. Exclusion did not vary by condition (10 per condition). One hundred and forty-eight (91 female) participants remained for the chips analyses.

Participants randomly assigned to the high cognitive effort condition (n = 80) completed a thought listing task during which they were forbidden to think of a white bear. Participants in the low cognitive effort condition (n = 68) also completed a thought listing task during which they were allowed to think of a white bear. Both conditions were given 6 minutes to complete the task. This manipulation has been used in past studies to vary cognitive effort (e.g., (54)). This experiment also included a second orthogonal manipulation to cognitive effort (room change) and several unrelated questions; these were not related to the hypotheses of the current research and will not be discussed further.

In experiment 1, after the cognitive task, participants were informed that the first experiment was over and they were starting the second experiment. Participants were given the opportunity to choose a flavor of potato chips (crisps) and were then given a canister containing 148g of chips. They were told that the study’s aim was to record students’ preferences for different types of chips. Participants were instructed to eat as many chips as they needed to accurately judge the taste, smell, and texture of the chips. Under this cover story, participants were asked to rate how much they enjoyed eating the chips using a 7-point scale (1 = not at all; 7 = very much enjoy). Because participants ate and thus rated a different number of chips, our key measure was the enjoyment rating after the first chip. Participants were given 10 minutes to complete the eating and rating task. After this time, the experimenter returned and weighed the chips cannister to determine chip consumption.

In experiment 2, 102 students at a Dutch university participated in exchange for partial course credit (in 2013). Five participants declined to consume the chocolate stimuli because of allergies. This did not differ by experimental condition (p = .198). Ninety-seven (53 female) participants remained for the chocolate analyses.

Participants randomly assigned to the high cognitive effort condition (n = 49) wrote about their daily routine without using words that contain the letters A or N. Participants in the low cognitive effort condition (n = 48) wrote the essay without using words that contained the letters X or Z. Both conditions were given 5 minutes to complete the task. Because there are more words than contain the letters A or N than X or Z, it requires more cognitive effort to override their natural way of writing. Participants completed a manipulation check “how much did you have to override or inhibit your typical way of writing in order to follow the instructions of the essay (1= Not at All; 7 = Very Much So)” to ensure the manipulation varied cognitive effort. The manipulation was successful: participants in the high (vs. low) cognitive effort condition reported overriding their natural way of writing more to complete the task [t(100) = 11.82, p < 0.001 (M = 6.17; SD = 1.48; M = 2.35, SD = 1.79)]. Thus our manipulation successfully varied cognitive effort during the essay task but it did not affect task compliance (i.e., how well they followed the essay instructions (1 = Very Poorly; 7 = Very Well); t < 1).

After the cognitive task, participants rated two types of stimuli presented in random order. To minimize carry-over effects and demand, these stimuli were presented as being ostensibly unrelated to the cognitive (writing) task. To test the effect of cognitive effort on desire for hedonically pleasurable objects, participants ate one piece of chocolate. Participants in this sample rated eating chocolate as a pleasant activity (M = 6.01, SD = 2.68), as evidenced by the rating differing significantly from the neutral midpoint of the scale, t(96) = 3.71, p < 0.001. After eating the piece of chocolate, participants indicated how yummy it was, how much they enjoyed eating it, how good it made them feel, and how much they wanted to eat more chocolate at that exact moment (0 = Not at all; 5 = Somewhat; 10 = Very much so). These items were averaged to form an index of enjoyment (α = 0.89).

To test the specificity of the effect, participants also rated the sensory qualities of four neutral items. Participants indicated the extent to which they perceived yellow post-it notes to be bright, a paper folder to be durable, a pen to be long, and the desk at which they were seated to be hard (0 = Not at all; 5 = Somewhat; 10 = Very much so). Not surprisingly, these items were not highly related (α = 0.50) so the rating for each stimulus was treated as a separate variable.

### Statistical analysis

For experiment in rats, data were analyzed in GraphPad Prism. Differences in cocaine taking and number of nose-poke responses were assessed by three-way ANOVA for repeated measures with session (1 to 22), drug (saline or cocaine) and cognitive exercise (effort or control) as factors. When the assumption of sphericity had been violated, data were analyzed using analysis of variance (ANOVA) corrected by the Greenhouse–Geisser method.

Differences in behavioral flexibility (measured as % of correct responses) were assessed by two-way ANOVA for repeated measures with session (1 to 22) and drug (saline or cocaine) as factors. Differences in the number of pellets obtained were assessed by two-way ANOVA with drug (saline or cocaine), cognitive exercise (effort or control) as factors. Differences in locomotor activity were assessed by one-way ANOVA with group (saline, Cocaine CE No Rest, Cocaine NoCE No Rest, Cocaine CE Rest, Cocaine NoCE Rest) as factor. Results showing significant overall changes were subjected to Tukey post-hoc test. Differences were considered significant when p<0.05.

For the experiment with humans, data were analyzed in SPSS. In experiment 1, we tested the effect of cognitive effort on enjoyment and consumption using an independent samples t-test. We then tested the indirect effect of cognitive effort on chip consumption through chip enjoyment using the Hayes Process Macro (v 3.5; Model 4). We tested the effect of cognitive effort on judgments using a repeated-measures ANOVA which treated the cognitive manipulation as a between-subjects factor and type of judgment (chocolate, post-it notes, paper, desk, pen) as a within-subjects factor. Since there was a significant interaction between the cognitive task and type of judgment, follow-up analyses were conducting using t-tests (i.e., the effect of the cognitive manipulation on judgments for each type of product). Differences were considered significant when p < 0.05.

Data are freely accessible via the Open Science Framework link: https://osf.io/7uhxf/

## Acknowledgments

This work was supported by the Centre National pour la Recherche Scientifique, the Institut National de la Santé et de la Recherche Médicale, the University of Poitiers, the Nouvelle Aquitaine CPER 2015-2020 / FEDER 2014-2020 program “Habisan” and the Nouvelle Aquitaine grant AAPR2020A-2019-8357510 (PI: M. Solinas) and the IRESP and the Aviesan Alliance grant « IRESP-19-ADDICTIONS-20 » (PI : M. Solinas).

This manuscript derives from experiments that were performed independently by the groups of M. Solinas and N. Mead and put together after discussion between M. Solinas and R. Baumeister in the course of a meeting organized by the CoCliCo team of the Cerca laboratory at the University of Poitiers.

We thank Michel Audiffren, Armand Chatard, Youna Vandaele and Nathalie Thiriet for helpful comments on a previous version of this manuscript.

This study has benefited from the facilities and expertise of PREBIOS platform (Université de Poitiers).

