## Supplementary Methods for "Cognitive effort increases the intensity of rewards"

Supporting Information

**Extended materials and methods**

**Experiments in rats**

**Subjects**

Seventy-five adult (10-11 weeks of age) male Sprague-Dawley rats (Janvier Labs and Charles Rivers, France) experimentally naive at the start of the study were used. Animals were individually housed in a temperature- and humidity-controlled room and maintained on a 12-hour light/dark cycle (light on at 7:00 AM) with free access to food and water. All experiments were conducted in accordance with European Union directives (2010/63/EU) for the care of laboratory animals and approved by the local ethics committees (COMETHEA).

**General Experimental Design**

Rats were pre-trained for 20 daily sessions on the set-shifting procedure in order to obtain a stable level of flexibility (Istin et al. 2017) before catheter implantation. After recovery, rats were divided into 4 experimental groups for each time conditions: cocaine cognitive effort (Coc CE), cocaine non-cognitive effort (Coc NoCE), saline cognitive effort (Sal CE), saline non-cognitive effort (Sal NoCE). For experiment 1, there was no interval between the cognitive session and the self-administration and the change from food to drug self-administration was associated simply by the switching off of the house light. For experiment 2, after the cognitive sessions, rats were removed from the self-administration cage and placed in their housing cage for a 2-4h period of rest before the self-administration session.

**Operant chambers**

Cognitive training and self-administration took place in the same operant chamber. Chambers were equipped with two levers on one wall that were used for the cognitive task and one single nose-poke on the opposite wall that was used for cocaine self-administration. A house light was used to discriminate the cognitive session (light on) from the self-administration session (light off). Diode lights above the levers were used for the attentional set-shifting task and sucrose pellets were delivered in a tray in the wall opposite to the levers and next to nose-poke. Inside the nose-poke there were red diode lights that were illuminated to indicate, together with the illumination of house light, the availability of cocaine. Responses on the nose-poke during the cognitive sessions and responses on the levers during the self-administration sessions were recorded but had no programmed consequences.

**Set-shifting task**

In a pilot experiment, we found that connection to the injection system during self-administration temporarily disrupts operant behavior for food. Therefore, to avoid this disruption, rats were implanted with a plastic screw in their back and were habituated to the connection to the metal spring used for self-administration during auto-shaping and pre-training.

Starting one week before the beginning of the experiments, food was restricted to approximately 15 – 18 g per day, which maintained rats at 85–90 percent of their weight. Feeding occurred in their home cages 1 hour after the experimental session. Rats had unlimited access to water.

Behavioral flexibility experiments were performed in Coulbourn experimental chambers controlled by GraphState software (Coulbourn Instruments, Allentown, PA, USA; www.coulbourn.com). The procedure was similar to the one described by Istin et al. (2017), with the difference that intertrial intervals lasted 15 sec instead than 1 sec. In brief, after shaping of operant responses, rats underwent 16 training sessions to assess trait behavioral flexibility. Each lever was surmounted by a diode green light that could be switched on or off according to the experimental phase of the procedure. The food tray was located on the opposite wall in order to force animals to reinitialize behavior and limit side biases. In each trial, in order to obtain food pellets, rats had to choose between the two levers based on two rules in different sensory dimensions: a visual dimension or an egocentric spatial dimension. For the visual rule dimension, animals had to follow the position of a light above one of the two levers, which indicated which lever was active in a given trial. For the egocentric rule dimension, animals had to keep responding on the same side (right or left) and to ignore the position of the light. In each session, if a rat produced 10 consecutive correct trials the rule was changed from one dimension to the other for a maximum of four different sets so that each dimension was presented twice per session. If animals did not reach this criterion, the trials continued with the same rule until the end of the experimental session. In each session there were four possible sets: (1) rule light (L); (2) rule side right (SR); (3) rule light (L) and (4) rule side left (SL). The session ended when a rat completed the four different sets or after 45 minutes whichever occurred first. Four configurations were possible: (1) L > SR > L > SL; (2) SR > L > SL > L; (3) L > SL > L > SR and (4) SL > L > SR > L. The starting set was counterbalanced among rats and changed daily for each rat so the four different configurations were presented every four sessions in ascending order. The number of total, correct and incorrect trials was measured. Flexibility was measured by the ability to adapt from one rule to the other, and therefore rats that showed lower percentage of errors were considered more flexible.

Rats assigned to the NoCE condition, underwent a procedure that was similar to the attentional set-shifting task except that every lever press during trials was reinforced with a pellet delivery regardless of position of the lever and whether the light was on or off. Thus, this group did not need to exert a cognitive effort to obtain rewards.

**Self-Administration Procedure**

Rats were allowed to self-administer cocaine or saline for sessions that lasted 150 minutes, according to Fixed Ratio (FR) schedule of reinforcement using a single nose-poke as operandum. The start of the self-administration session was signaled only by the switching off of the house light and intermittent illumination of the nose-poke. Initially, FR value was set at 1 and, after 7 sessions, it was increased to 3. During the self-administration, completion of the FR resulted in the immediate delivery of an i.v. injection of cocaine (0.15mg/inj) or saline (0.9% NaCl). Following the i.v. injection, the nose-poke cue light pulsed for 10 seconds and was followed by 20 seconds time-out indicated by the illumination of the houselight. During the time-out period, responding on the nose-poke was recorded but had no programmed consequences.

**Locomotor activity experiments**

Separate groups of rats were trained for 20 sessions in the set-shifting task. After this training, half of the rats continued to perform the cognitive task (cognitive effort, CE) to obtain food pellets whether the other half, were reinforced regardless of the rule (No Cognitive effort, NoCE) for 5 additional sessions. After the last session, rats were injected with cocaine (10 mg/kg i.p.) either immediately or 3h after the cognitive session and were put in an open field where locomotor activity was recorded by a viewpoint videotracking system (<https://www.viewpoint.fr>) for 1h.
