## Supplementary material for "Cognitive effort increases the intensity of rewards": Fig.S1-5

### Slide 1
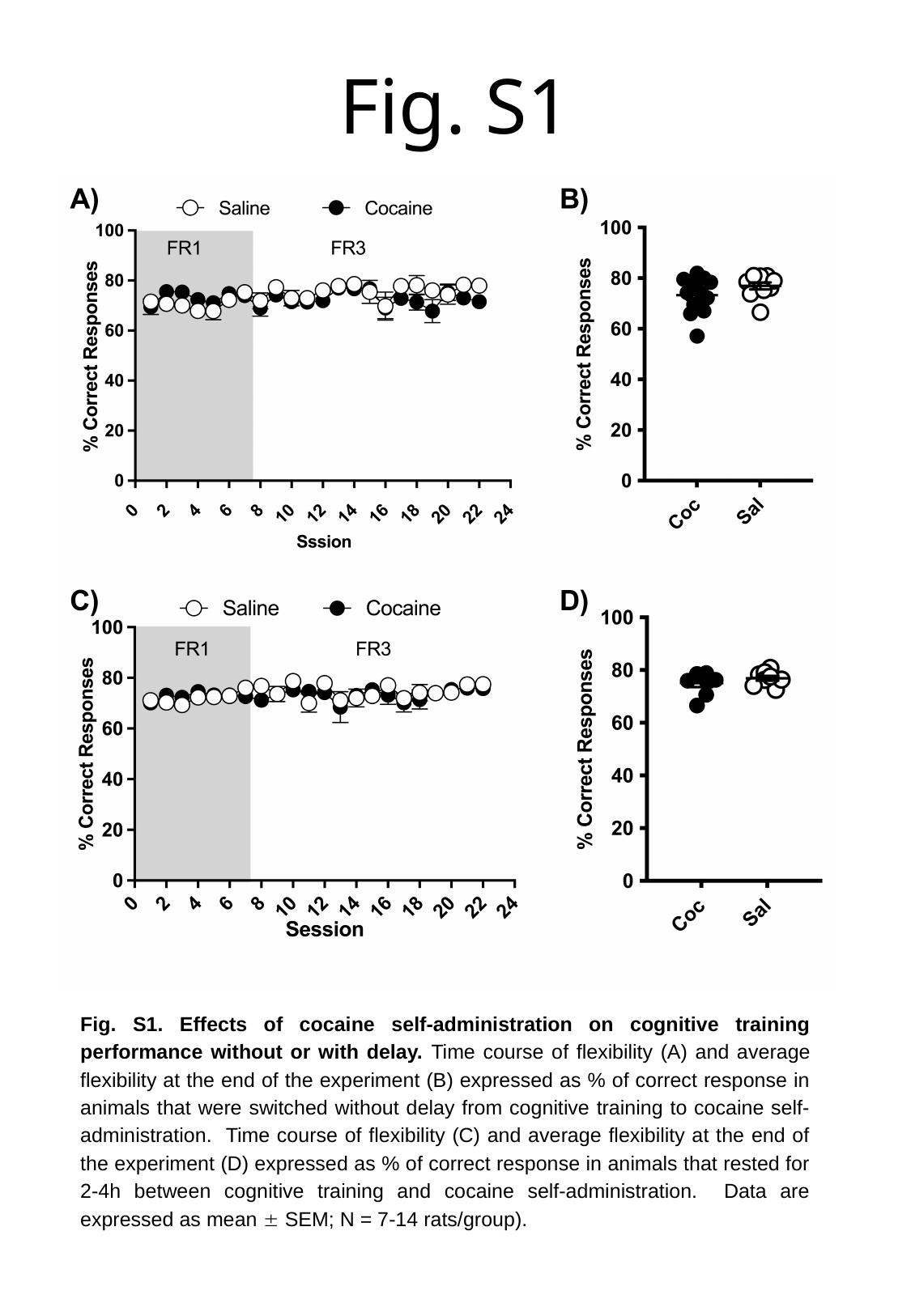

#### Fig. S1
Fig. S1. Effects of cocaine self-administration on cognitive training performance without or with delay. Time course of flexibility (A) and average flexibility at the end of the experiment (B) expressed as % of correct response in animals that were switched without delay from cognitive training to cocaine self-administration. Time course of flexibility (C) and average flexibility at the end of the experiment (D) expressed as % of correct response in animals that rested for 2-4h between cognitive training and cocaine self-administration. Data are expressed as mean ± SEM; N = 7-14 rats/group).

### Slide 2
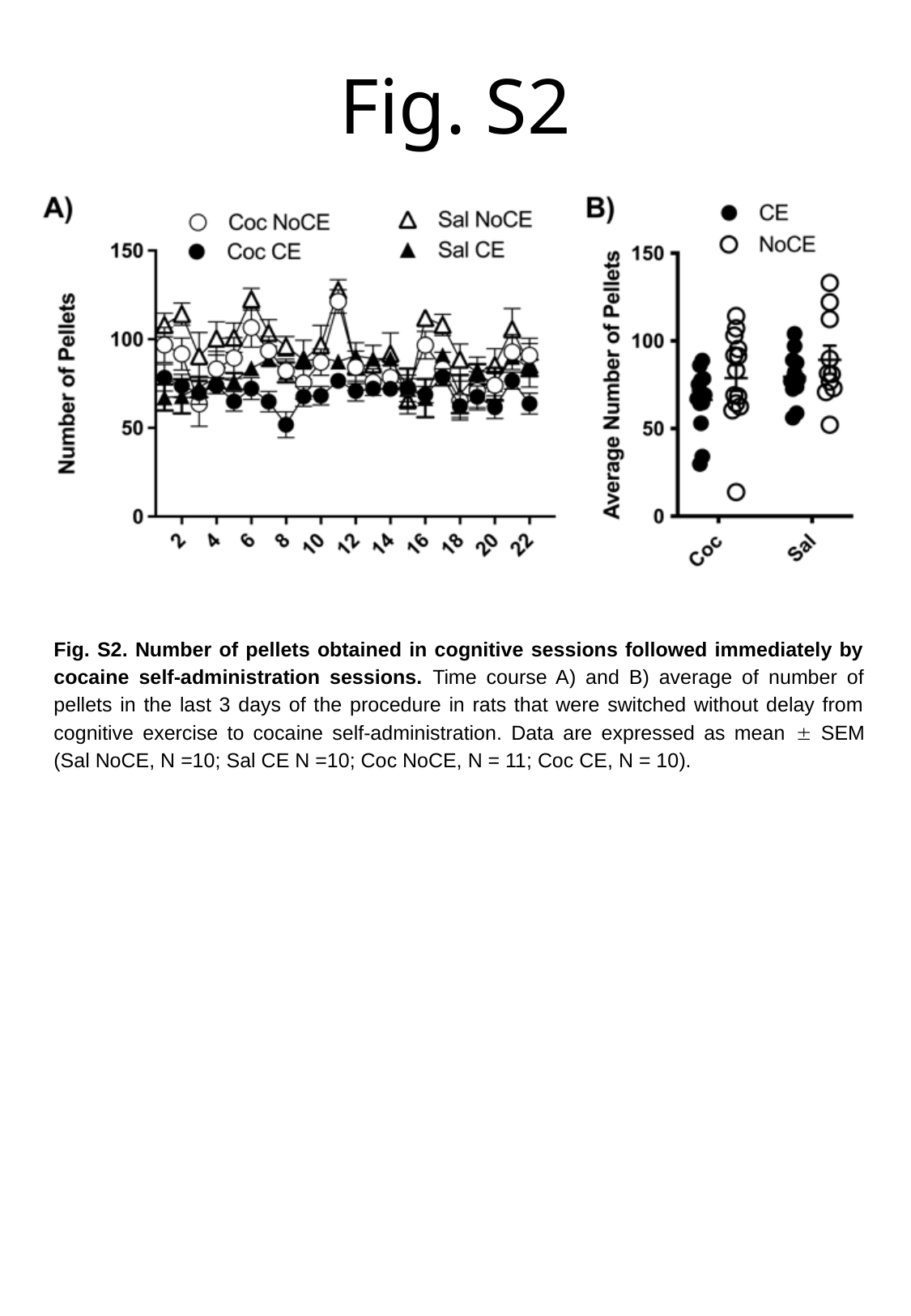

#### Fig. S2
Fig. S2. Number of pellets obtained in cognitive sessions followed immediately by cocaine self-administration sessions. Time course A) and B) average of number of pellets in the last 3 days of the procedure in rats that were switched without delay from cognitive exercise to cocaine self-administration. Data are expressed as mean ± SEM (Sal NoCE, N =10; Sal CE N =10; Coc NoCE, N = 11; Coc CE, N = 10).

### Slide 3
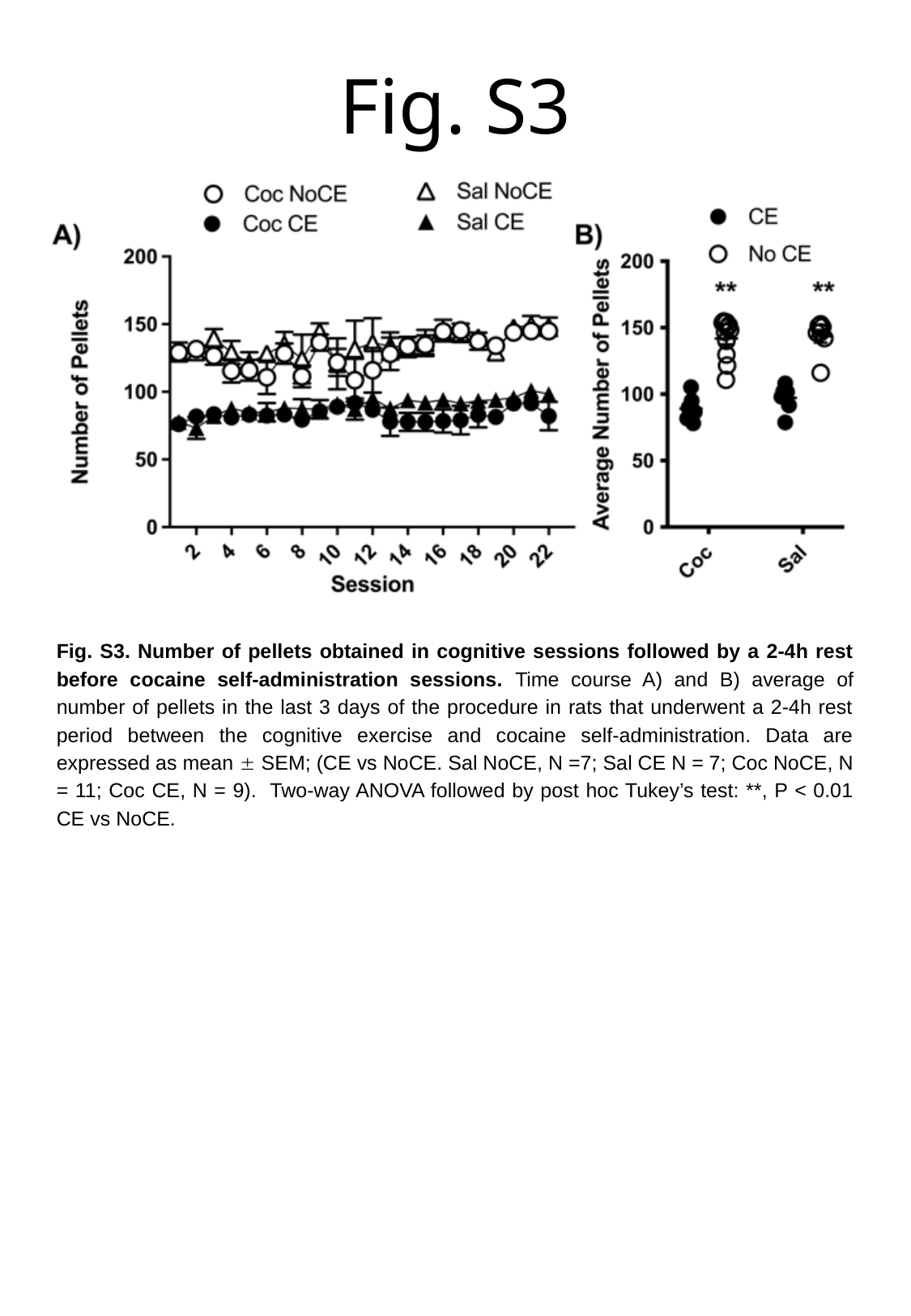

#### Fig. S3
Fig. S3. Number of pellets obtained in cognitive sessions followed by a 2-4h rest before cocaine self-administration sessions. Time course A) and B) average of number of pellets in the last 3 days of the procedure in rats that underwent a 2-4h rest period between the cognitive exercise and cocaine self-administration. Data are expressed as mean ± SEM; (CE vs NoCE. Sal NoCE, N =7; Sal CE N = 7; Coc NoCE, N = 11; Coc CE, N = 9). Two-way ANOVA followed by post hoc Tukey’s test: **, P < 0.01 CE vs NoCE.

### Slide 4
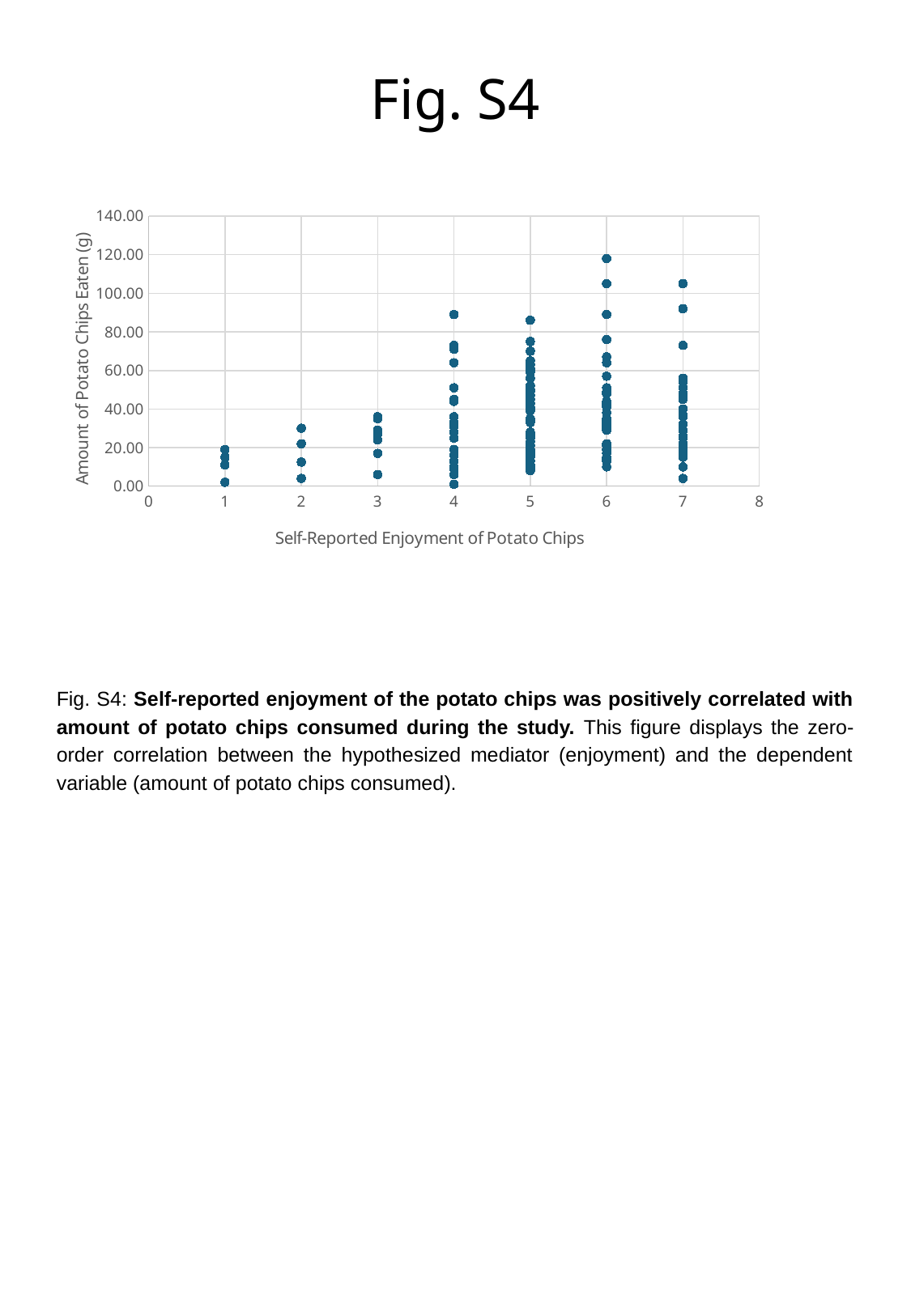

Fig. S4
###### Chart:
| Category | |
|---|---|Fig. S4: Self-reported enjoyment of the potato chips was positively correlated with amount of potato chips consumed during the study. This figure displays the zero-order correlation between the hypothesized mediator (enjoyment) and the dependent variable (amount of potato chips consumed).

### Slide 5
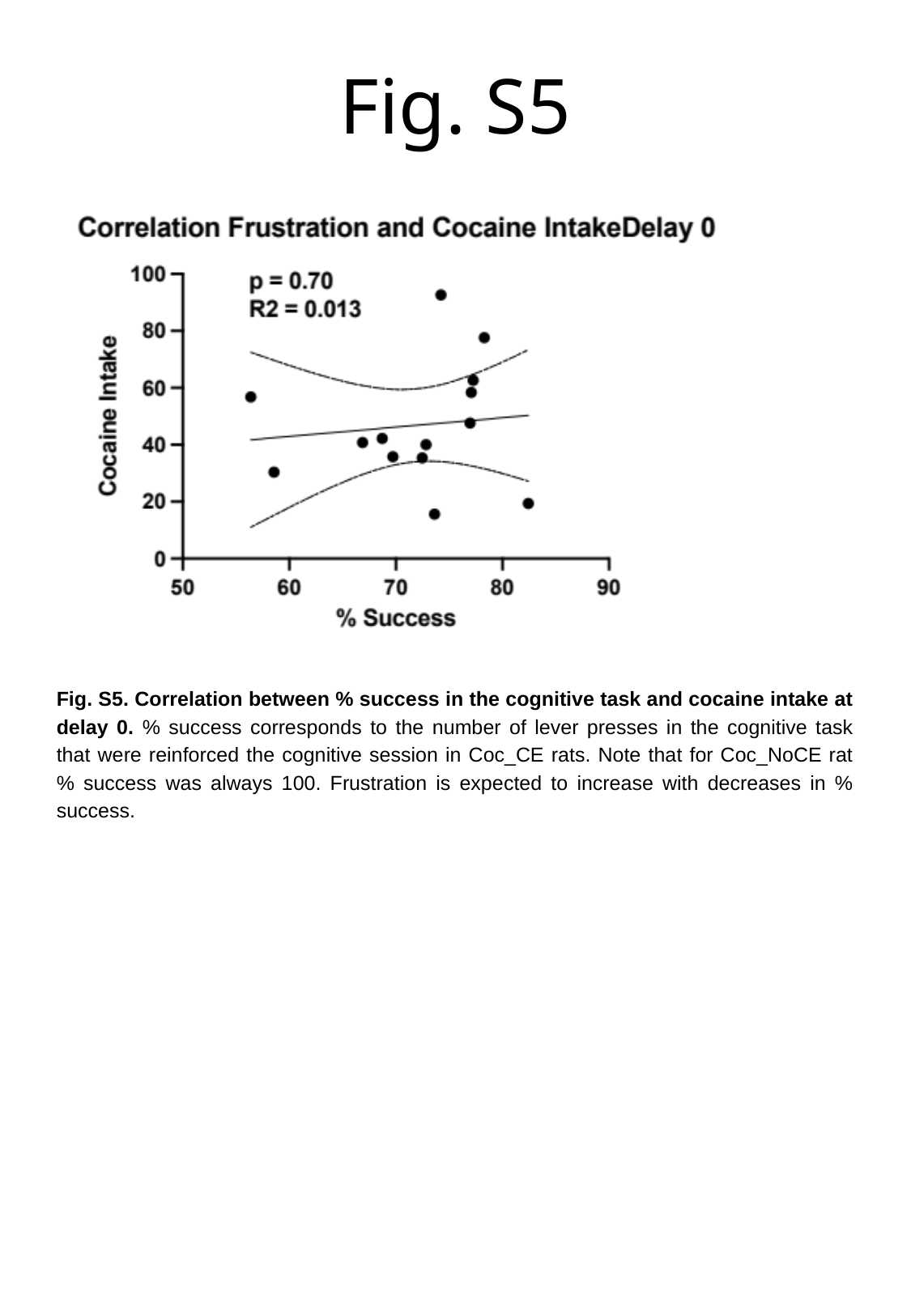

#### Fig. S5
Fig. S5. Correlation between % success in the cognitive task and cocaine intake at delay 0. % success corresponds to the number of lever presses in the cognitive task that were reinforced the cognitive session in Coc_CE rats. Note that for Coc_NoCE rat % success was always 100. Frustration is expected to increase with decreases in % success.
